# Excitation-Inhibition Balance and Fronto-Limbic Connectivity Drive TMS Treatment Outcomes in Refractory Depression

**DOI:** 10.1101/2025.04.23.648963

**Authors:** Davide Momi, Zheng Wang, Mohammad P. Oveisi, Kevin Kadak, Sorenza P. Bastiaens, Jennifer I. Lissemore, Yoshihiro Noda, Jonathan Downar, Fidel Vila Rodriguez, Rebecca Strafella, Zafiris J Daskalakis, Christoph Zrenner, Reza Zomorrodi, Camarin E. Rolle, Manish Saggar, Nolan Williams, Corey J. Keller, Daniel M. Blumberger, Daphne Voineskos, John D. Griffiths

## Abstract

Depression affects over 350 million people worldwide, with treatment resistance occurring in up to 30% of cases. Intermittent theta burst stimulation (iTBS) targeting the left dorsolateral prefrontal cortex (DLPFC) has emerged as a promising intervention, yet the neurophysiological mechanisms determining which patients will respond remain poorly understood. Here, we combined transcranial magnetic stimulation with electroencephalography and whole-brain computational modeling to uncover the mechanistic basis of treatment efficacy in 90 patients with treatment-resistant depression. We identified two distinct neurophysiological signatures that differentiate responders from non-responders: (1) post-treatment shifts in excitation-inhibition balance toward greater inhibitory control, and (2) a pre-treatment brain state characterized by anticorrelated dynamics between subgenual anterior cingulate cortex and DLPFC. These features were significantly correlated with clinical improvement and could not be explained by non-specific factors. Our findings provide a neurophysiologically-informed framework for developing personalized and optimized neuromodulation approaches in treatment-resistant depression.

## MAIN

Major depressive disorder (MDD) has a lifetime prevalence of approximately 20% in the US and affects over 350 million people worldwide, making it one of the leading contributors to disability^1^. Despite various available treatments, only about 30% of patients achieve remission with first-line therapies^2^, and many develop treatment-resistant depression—a condition where patients fail to respond to multiple medication trials and face increasingly limited therapeutic options. Repetitive transcranial magnetic stimulation (rTMS) has emerged as a promising intervention for these patients with TRD^3–5^, offering an alternative where conventional pharmacotherapy has failed.

Intermittent theta burst stimulation (iTBS), an FDA-approved rTMS protocol that couples slow theta-frequency carrier waves with brief gamma-frequency bursts, has garnered particular attention for its enhanced neurophysiological effects and treatment efficiency compared to conventional rTMS^6^. iTBS delivered to the left dorsolateral prefrontal cortex (L-DLPFC) has demonstrated promising response and remission rates in TRD^7^. Yet despite these advances, treatment outcomes remain highly variable, with response rates typically ranging from 29% to 46%^8,9^. This variability in treatment response presents a significant clinical challenge, and underscores a fundamental gap in our understanding of depression pathophysiology and neuromodulation mechanisms.

To address this clinical variability and advance therapeutic outcomes, we must bridge the gap between basic neuroscience and clinical application. Specifically, two critical questions emerge that lie at the intersection of basic neuroscience and clinical psychiatry: i) What are the factors that determine whether a given administration of iTBS therapy will effectively engage the neural circuit and neuroplasticity-related mechanisms that ultimately lead to alleviation of depressive symptoms in a given patient? ii) What neurophysiological markers can be used in the clinic to reliably distinguish between potential responders and non-responders before treatment begins, and what can we say about the neural circuitry underlying these markers? Answering these could transform the field of TRD neuromodulation therapy design, moving away from its current paradigm of trial-and-error discovery science, and moving toward a more effective precision medicine model, where interventions are tailored to the specific individual features of the patient’s own neuroanatomy and neurophysiology. Multiple neuroimaging approaches have sought to answer these two questions over the past 20 years, within which TMS-EEG stands out as the most powerful tool available for flexibly studying the neural mechanisms underlying iTBS treatment effects. By combining TMS with EEG, researchers can measure cortical excitability and plasticity through both stimulation-evoked responses and stimulation-induced oscillatory activity. This approach has identified neurophysiological markers that differentiate responders from non-responders across various TMS protocols^10–13^, with particular emphasis on treatment-related changes in low-frequency oscillations, that may reflect fundamental alterations in cortical circuit properties. Specific components of the TMS-evoked potential (TEP) and induced oscillatory activity serve as indirect proxies of cortical excitation-inhibition (E/I) balance^14,15^, which appears disrupted in MDD^16^. Indeed, Voineskos et al.^17^ demonstrated significant deficits in GABA-mediated cortical inhibition in patients with MDD compared to healthy controls, aligning with broader evidence of altered GABA and glutamate levels across multiple brain regions in depression^18–20^. Notably, better clinical outcomes have been associated with specific pre-treatment TMS-EEG markers, including a more negative N45 waveform, and a smaller P60 amplitude^21^, potentially reflecting enhanced GABAergic inhibition and reduced glutamatergic excitation, respectively.

Complementing these electrophysiological findings, functional magnetic resonance imaging (fMRI) studies have revealed that stronger negative functional connectivity between the DLPFC stimulation site and subgenual anterior cingulate cortex (sgACC) at baseline predicts better treatment outcomes^22–25^. This latter finding highlights the importance of fronto-limbic circuit dynamics in determining treatment response, suggesting that rTMS may achieve its therapeutic effects by modulating communication between these key regions.

Despite these promising leads however, the field of therapeutic TMS does face significant challenges. Results show high inter- and intra-subject variability, findings across studies often appear contradictory, and the precise mechanisms linking observed neural responses to clinical outcomes remain elusive. This knowledge gap severely hampers efforts to optimize and personalize TMS treatments. Neuroimaging and human neurophysiology techniques for measuring and monitoring brain activity, while valuable, allow only indirect inferences about underlying neural mechanisms, and carry a host of limitations such as low signal-to-noise ratios, limited spatial resolution, and challenges in accurate measurement^26,27^. What is critically needed is an integrative mechanistic framework that can bridge between measurable brain signals and the underlying neural circuit dynamics that determine treatment response. Whole-Brain Modeling (WBM) - the sub-field of computational neuroscience concerned with the theoretical principles and numerical simulation of large-scale brain network dynamics - offers just such a framework. WBMs can provide a robust, flexible approach for testing mechanistic hypotheses about how iTBS modulates neural circuit dynamics to achieve therapeutic effects^28,29^. This approach has already demonstrated value in understanding the pathophysiology of various conditions^30–34^.

In this study, we combined longitudinal TMS-EEG measurements with WBM to investigate the neurophysiological mechanisms underlying treatment response in TRD. Our integrated approach revealed two key findings: First, successful iTBS treatment was characterized by specific reductions in low-frequency (3–10 Hz) oscillatory power. Computational modeling indicated that these power changes reflected shifts in E/I balance toward greater inhibitory control, resulting from reduced excitatory drive to cortical pyramidal cell populations. Second, a specific spatiotemporal activity pattern observed in pre-treatment TMS-EEG, characterized by an anti-phase (negatively correlated) relationship between left DLPFC and sgACC, was found to predict clinical outcomes - consistent with fMRI studies of fronto-limbic connectivity in TRD^22–24^. Together, these findings provide a neurophysiologically-grounded framework linking treatment efficacy to fundamental changes in cortical dynamics, pointing toward novel approaches for personalizing neuromodulation therapies in TRD.

## RESULTS

### Higher inhibition in responders following iTBS

We first examined the differential impact of iTBS on the induced oscillatory activity of stimulus-related brain dynamics. To do this, we compared TMS-EEG time-frequency activity between responders and non-responders. As shown in Fig. 1A, we compared the post-iTBS minus pre-iTBS induced power difference between responders and non-responders. While both groups showed decreases in induced power following iTBS, responders exhibited significantly greater reduction in the 3-10 Hz range compared to non-responders. This differential spectral power response was statistically significant (1,000 permutations, all clusters p < 0.05, corrected). The significant cluster spanned the 3-10 Hz frequency range and was most pronounced between 50-200ms post-stimulus, indicating a distinct neurophysiological response pattern in participants who responded to the iTBS intervention versus those who did not. Transient differences in gamma-band activity (30–50 Hz) were also observed, but these effects did not survive cluster-based permutation testing. To better understand the circuit mechanisms underlying this low-frequency TEP suppression effect, we next turned to the WBM approach outlined above (see also Fig. 5) - first with data-driven physiological parameters estimated from model fits to TEP waveform data, followed by some novel exploratory numerical simulations.

**Fig. 1.**
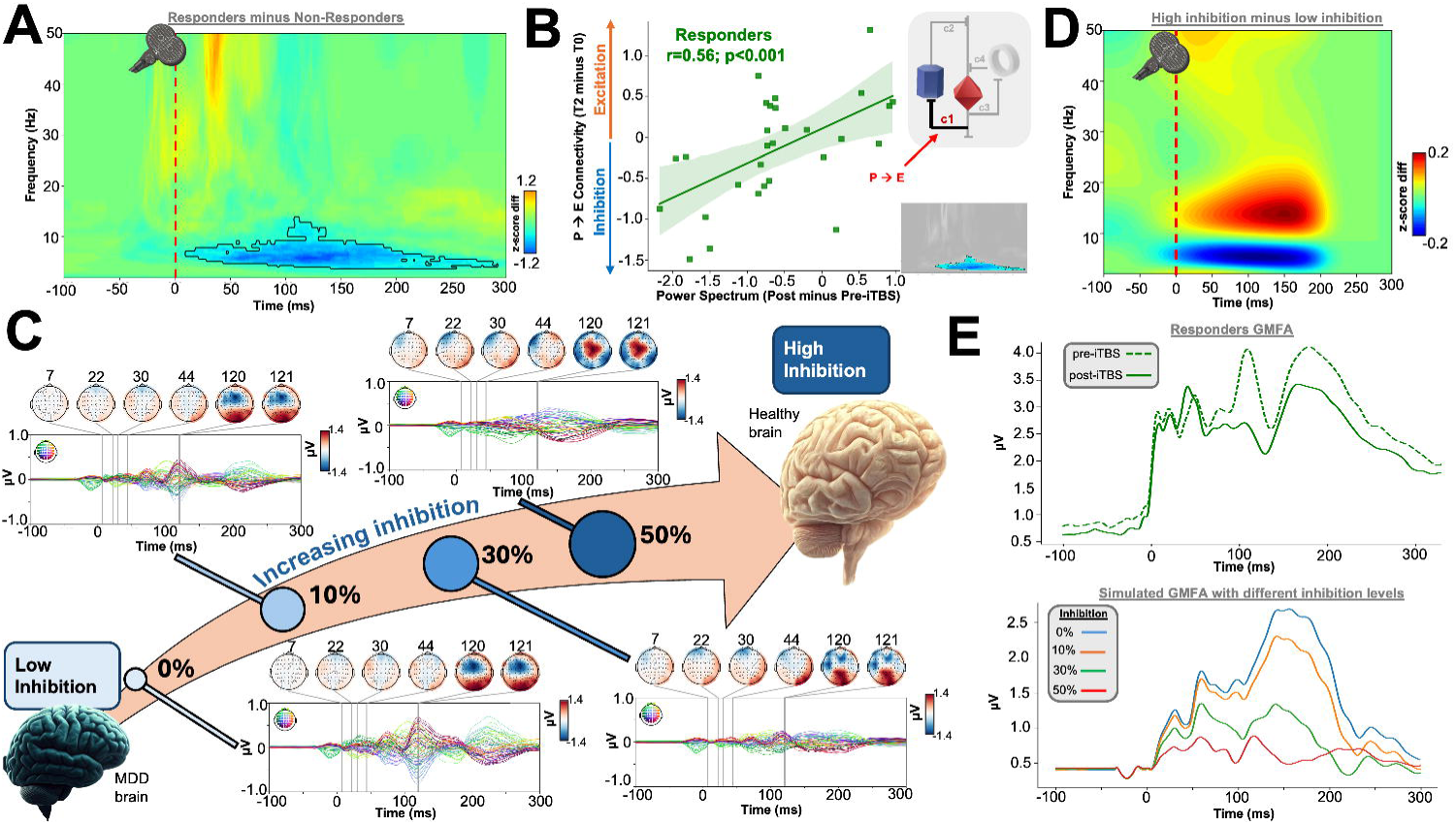
iTBS therapy suppresses TEP low-frequency oscillatory power by modulating the level of inhibition. **(A)** Induced time-frequency spectral response difference between responders and non-responders, showing significantly lower power in the 3–10 Hz range in responders relative to non-responders. **(B)** Significant correlation (r = 0.56, p < 0.001) between post-iTBS vs pre-iTBS changes in the P→E synaptic weights) and in the significant cluster regions identified in Panel A. **(C)** Simulated effects of shifting E/I balance toward inhibition (via decreased P→E) for the participant with the highest Hamilton Depression Rating Scale score (indicating more severe MDD). This simulation reveals a notable reduction in TEP amplitude, especially in the late components, with increasing inhibition. **(D)** Group-level induced time-frequency spectral comparison between the original (0% increased inhibition) and a 50% inhibition-increased simulation, demonstrating a similar low-frequency reduction as observed in empirical data in Panel A. **(E)** GMFA for empirical data, with pre (dashed) vs. post (solid) responses shown for responders (top). GMFA across simulations with incremental inhibition (bottom) reveals a similar trend where increased inhibition is associated with a decrease in the late component amplitude for responders.

Our model of neurostimulation-evoked brain dynamics^35,36^ is a connectome-based extension of the classic Jansen-Rit (JR) neural mass model^37–39^. In JR, shifts in the excitatory/inhibitory (E/I) balance can occur through complementary synaptic mechanisms: decreased pyramidal drive to excitatory cells (reduced P→E; c1) and enhanced inhibitory feedback (increased I→P; c4). Both mechanisms functionally shift the circuit toward greater inhibitory influence, though through distinct synaptic pathways. Throughout the following, for the sake of clarity we refer to both of these related changes as shifts toward inhibition.

Looking at the model parameter estimates in relation to the TEP waveform properties: a significant positive correlation was found between changes (post-iTBS minus pre-iTBS) in the excitatory feedback loop strength (expressed as P→E cell connection strength, c1) and the z-score differences in 3–10 Hz power between post-iTBS and pre-iTBS observed in responders (Fig. 1B; r = 0.56, p < 0.001). This correlation indicates that a greater reduction in low-frequency power, as observed in responders to iTBS treatment (Fig. 1A), is associated with reduced excitatory feedback after treatment. This result suggests that the plasticity-inducing effects of iTBS may decrease excitatory feedback mechanisms, leading to the observed diminishment in low-frequency oscillatory responses.

To further explore the mechanistic role of increased inhibition, we simulated the effects of reducing the P→E cell connection strength by 10%, 30%, and 50% of the original value across all participants. For illustration, Fig. 1C presents results of this simulation on the time-domain TEP waveforms for the participant with the highest Hamilton Depression Rating Scale (HDRS) score^40^, representing more severe MDD. This simulation, which shifted the circuit balance toward inhibition by decreasing the P→E connection strength, resulted in a marked reduction in the TEP amplitude, particularly in the late components. The impact of enhanced inhibitory signaling appears to attenuate TEP responses in patterns consistent with those observed in clinical responders.

Corresponding analyses in the time-frequency domain are shown in Fig. 1D, where we compared simulated TMS-EEG data with baseline (0% increase) and enhanced (50% increase) levels of inhibition. Similar to the empirical findings in responders (Fig. 1A), simulations showed a reduction in low-frequency power when inhibition was elevated, aligning with the 3–10 Hz power reduction observed in Panel A.

Finally, we examined the Global Mean Field Amplitude (GMFA)^41^ across empirical and simulated conditions (Fig. 1E). In the empirical data, responders showed a decrease in GMFA post-iTBS in the late component of the evoked response (solid line) compared to the pre-iTBS baseline (dashed line), replicating our previous result with this dataset^42^. Simulations with incremental inhibition showed a similar trend, where increasing inhibition progressively reduced the GMFA, particularly in the late component for responders. These converging lines of evidence further strengthen the relationship between iTBS-induced inhibitory modulation and treatment response^43^.

### Responder-specific elevated inhibition after iTBS treatment

To examine differential neural responses to iTBS, we analyzed changes in JR model parameters between responders and non-responders. The 2×2 repeated measures Analysis of Variance (ANOVA) for inhibitory-to-pyramidal (I→P; c4) synaptic weights revealed a significant interaction time*group (F(1,86) = 29.11, p < 0.001) and a significant main effect of group (F(1,86) = 76.86, p < 0.001). Post-hoc tests showed no pre-iTBS differences (t = -1.21; p = 0.23) but significant post-iTBS differences between groups (t = -2.598; p=0.0147), with responders exhibiting greater I→P weights (Fig. 2A). As I→P was the only parameter showing significant group effects, we further explored its predictive value by correlating pre-iTBS I→P weights with post-treatment HDRS scores. This analysis revealed a significant negative correlation (r = - 0.56, p < 0.0001), indicating that higher inhibitory feedback pre-treatment predicted greater symptom improvement among responders (Fig. 2B).

**Fig. 2.**
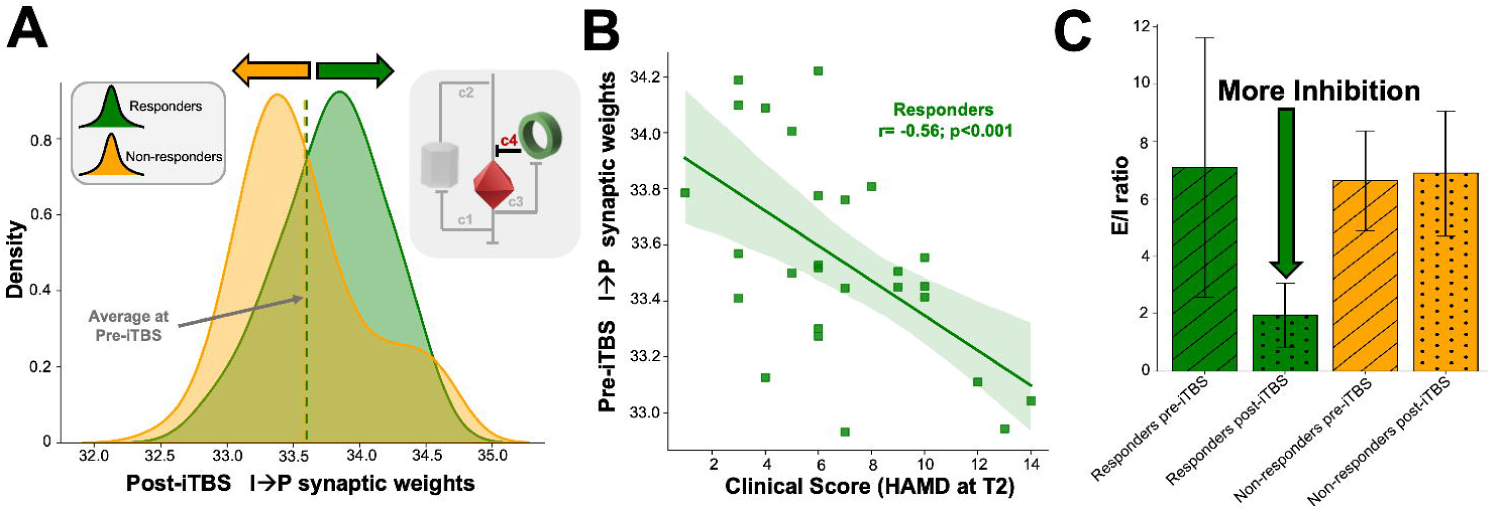
iTBS therapy modifies inhibitory feedback in cortical circuits differently in responders and non-responders. **(A)** Strength of the inhibitory feedback loop for responders (green) and non-responders (yellow) post-iTBS therapy, as given by per-group distributions of estimated Inhibitory interneuron (I) to Pyramidal cell (P) population synaptic weights (I→P). Vertical dashed line indicates the average pre-iTBS value of this parameter, showing that both responders and non-responders had the same starting value before treatment. Post-iTBS therapy, responders exhibit an increase in inhibitory feedback, while non-responders show the opposite effect. **(B)** Pre-iTBS I→P synaptic weights demonstrate a significant negative correlation with post-treatment depression severity (HDRS scores) exclusively in responders (r = -0.56, p < 0.0001). This relationship indicates that patients with higher inhibitory feedback prior to treatment experienced greater symptom reduction, suggesting that iTBS may exert its therapeutic effects by enhancing inhibitory mechanisms within local neural circuits **(C)** Ratios of excitatory to inhibitory (E/I) synaptic weights across responders (green) and non-responders (yellow), both before (dashed) and after (dotted) iTBS therapy. A significant increase in inhibition is observed in responders only following the treatment.

For E/I balance, the ANOVA also revealed a significant group*time interaction (F(1,86) = 38.29, p < 0.001) and main effect of group (F(1,86) = 59.21, p < 0.001). While pre-iTBS E/I balance showed no between-group differences (t = 0.82, p = 0.19), post-iTBS measures differed significantly (t = 10.088, p<0.001). This shift suggests that iTBS selectively modulated inhibitory mechanisms in responders, while non-responders showed no significant E/I balance changes (Fig. 2C). These findings align with literature suggesting increased inhibitory tone may benefit treatment outcomes in responders^10,44^.

### Subgenual-prefrontal interaction predicts the efficacy of iTBS therapy

Our results reported thus far point to modulation of inhibition, of E/I ratios, and of low-frequency TMS-induced oscillatory power as signatures of successful rTMS therapy for TRD, One limitation of these findings however is that they both represent fairly global and spatially non-specific markers of both brain activity (GMFA) and brain circuit physiology (P→E, I→P). Next, we explored whether our patient-personalized WBMs of TMS-evoked brain dynamics contained meaningful spatiotemporal patterns of neural activity, and whether these showed any relationship to clinical improvement.

Mean-centered Partial Least Squares (PLS) analysis on the model’s pre-treatment neural population time series identified a significant brain state (Fig. 3A) that maximally differentiates between responders and non-responders (p=0.001). Interestingly, within this brain state, the eigenvector values extracted from the left DLPFC (the area stimulated during iTBS) and left sgACC (a deeper mesocortical limbic system region strongly implicated in MDD pathophysiology) exhibited opposite signs, as indicated by their respectively red and blue colours in Fig. 3A/B. The opposite-polarity loadings of DLPFC and sgACC on this treatment-related eigenvector align with the extensive prior fMRI literature reporting that the strength of (resting-state BOLD time series) anti-correlations between sgACC and DLPFC rTMS target location predicts therapeutic outcomes^22–24,45^.

**Fig. 3.**
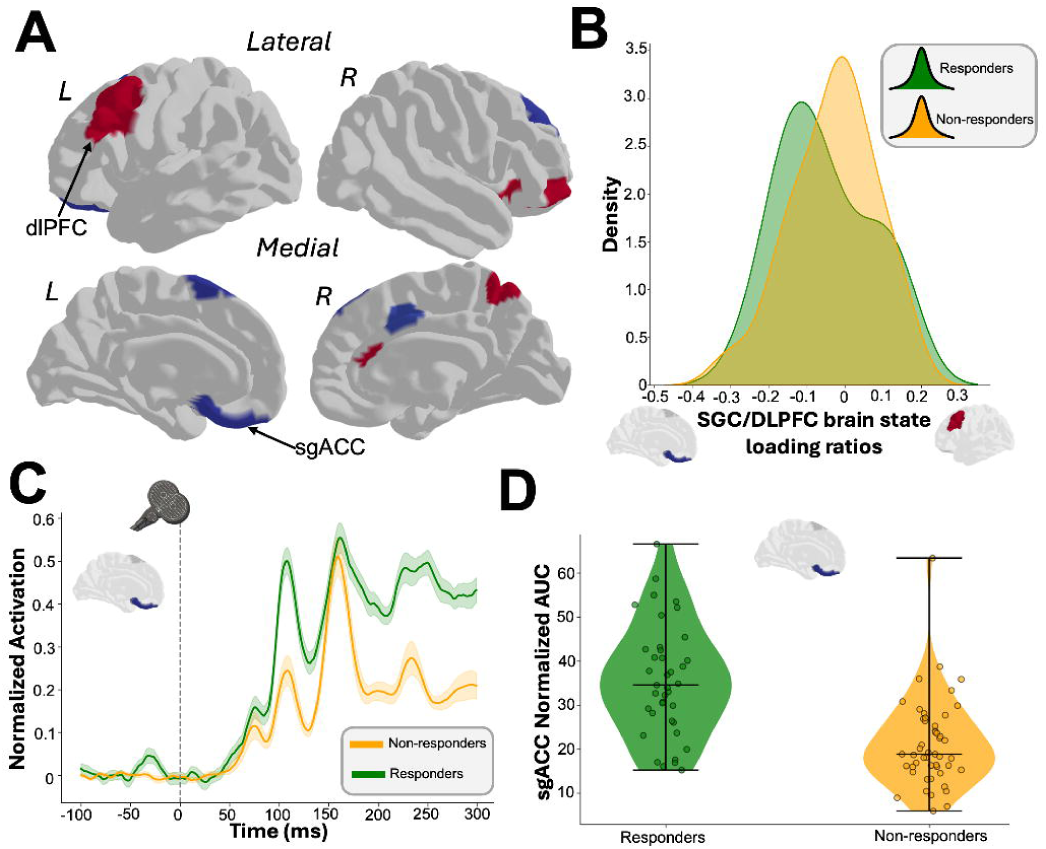
Neural Signatures Predict Clinical Response to iTBS Treatment for Depression. **(A)** Brain state that maximizes the difference between responders and non-responders following iTBS treatment. Lateral and medial views highlight the dlPFC and sgACC regions showing opposite eigenvector loading directions. **(B)** Ratio of sgACC/DLPFC loadings within the brain state maps in A) for responders (green) and non-responders (yellow), showing a negative relationship for responders only. **(C)** Normalized sgACC activation measured by GMFA at pre-iTBS baseline for responders (green) and non-responders (yellow). Time courses demonstrate higher sgACC engagement patterns for responders. **(D)** AUC analysis of sgACC activation for the entire cohort, demonstrating significantly higher normalized engagement in responders compared to non-responders (p<0.001), suggesting pre-treatment sgACC activity may serve as a predictive biomarker.

Having identified this TEP model-based brain state signature as showing sensitivity to treatment efficacy, we next asked whether it was expressed differentially in different outcome-defined patient groups in their baseline, pre-treatment brain activity. This was indeed found to be the case (Fig. 3B), with responders showing a negative relationship between sgACC and DLPFC in their brain state loadings at baseline, whereas non-responders did not exhibit this pattern.

Further investigation of the temporal dynamics of sgACC activation revealed distinct patterns between responders and non-responders (Fig. 3C). Time course analysis demonstrated significantly higher sgACC engagement patterns in responders (green) compared to non-responders (orange) following pre-iTBS baseline stimulation. These temporal differences in sgACC recruitment suggest fundamental differences in cortico-limbic circuit dynamics that may underlie treatment susceptibility.

To quantify the difference in sgACC engagement more precisely, we conducted Area Under the Curve (AUC) analysis of sgACC activation across the entire cohort (Fig. 3D). This analysis revealed significantly higher normalized engagement in responders compared to non-responders (p<0.001), with responders showing a mean normalized AUC of 35.6 ± 4.2 compared to 21.8 ± 3.8 in non-responders. The clear separation between the groups in pre-treatment sgACC activity supports its potential value as a predictive biomarker for iTBS treatment outcomes in depression. This finding aligns with the growing evidence that baseline neurophysiological states, particularly involving sgACC connectivity, may determine therapeutic responsiveness to neuromodulation interventions^23^.

Pre-treatment values of this prefrontal-subgenual corticolimbic circuit could, for example, provide a novel biomarker for predicting treatment outcomes.

### Greater trajectory deviation and increased stability in neural dynamics of iTBS responders

One of the advantages of our WBM-based approach to analyzing TMS-EEG data is that it allows us to interpret TMS-evoked potentials both in their conventional time-series representation and as geometric trajectories within the system’s phase space (Fig. 4A). The latter allows certain dynamical features such as attractor manifolds and recurrent trajectories to be visualized and characterized. We quantified the iTBS treatment-induced modifications to this attractor landscape by computing point-wise Euclidean distances in the treatment-predictive brain state between post-iTBS and pre-iTBS trajectories for responders and non-responders (Fig. 4B). A group comparison using non-parametric permutation testing (10,000 iterations) revealed that responders exhibited significantly greater trajectory deviation following iTBS compared to non-responders (t = 3.24, p < 0.01), indicating a more substantial shift in the underlying dynamical regime induced by the intervention.We also examined the distance of each group’s post-iTBS trajectory from the stable fixed point of the system (Fig. 4C). Responders remained significantly closer to the attractor (t = 2.18, p < 0.05), suggesting increased stability and reduced susceptibility to external perturbations, such as TMS pulses. Together, these results provide evidence that iTBS induces a meaningful reconfiguration of brain state dynamics in responders, characterized by both greater reorganization (Fig. 4B) and increased attractor convergence (Fig. 4C). For dynamic visualizations of the state-space trajectories, see supplementary videos V1 and V2.

**Fig. 4.**
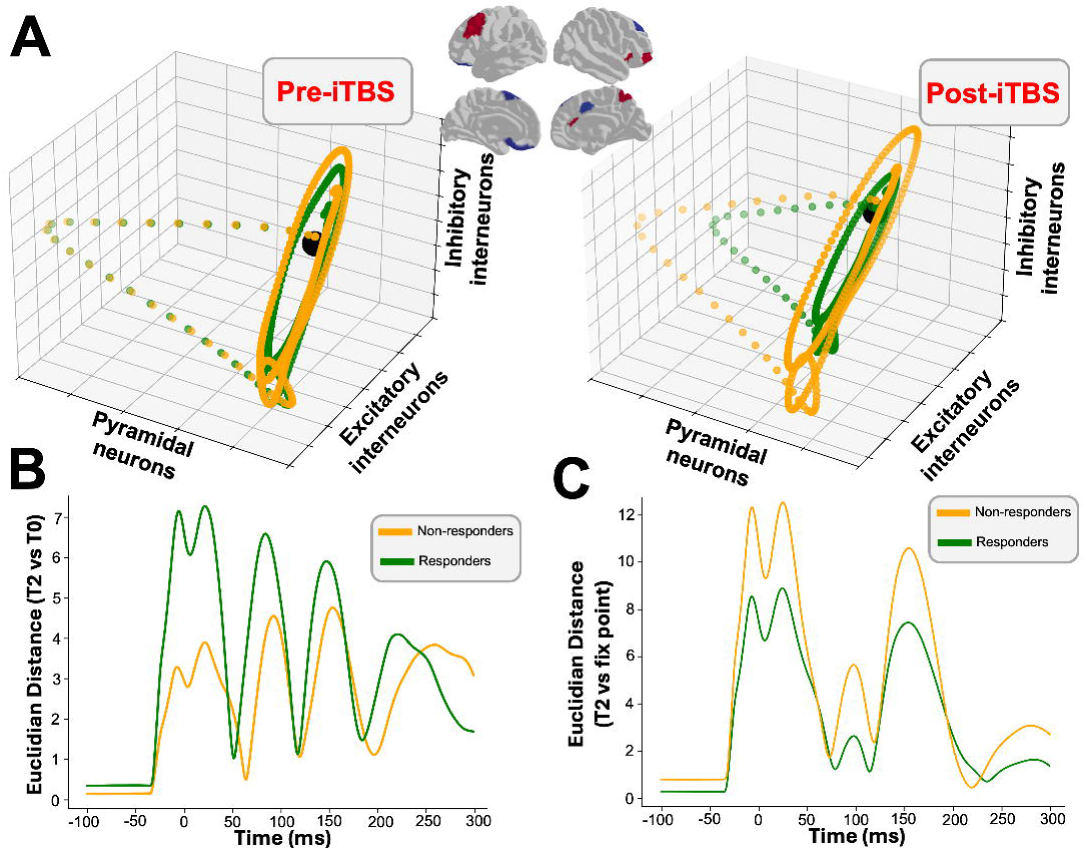
iTBS selectively reorganizes attractor dynamics in treatment-responsive individuals. **(A)** Three-dimensional neural trajectories of the treatment-predictive brain state (as identified in Fig. 3 for responders (green) and non-responders (yellow), shown pre-iTBS (left) and post-iTBS (right). Pre-treatment, both groups exhibit similar trajectories, remaining near the stable fixed point (black dot). Post-treatment, responders show a marked trajectory contraction and reduced deviation from the fixed point, suggesting greater stability and reduced sensitivity to external stimuli (e.g., TMS). **(B)** Point-wise Euclidean distances between post-iTBS and pre-iTBS trajectories. Responders display significantly greater trajectory shifts, indicating a more substantial reconfiguration of the treatment-predictive brain state following iTBS. **(C)** Point-wise Euclidean distance from the stable fixed point in the post-iTBS condition. Responders remain consistently closer to the fixed point, suggesting reduced susceptibility to external perturbation and enhanced attractor stability compared to non-responders.

## DISCUSSION

Our investigation offers several novel insights into the fundamental neurophysiological mechanisms underlying successful iTBS therapy in treatment-resistant depression, arrived at through a novel computational framework for personalized brain network modeling and neuroimaging data analysis^35,36^. In particular, the integration of empirical TMS-EEG measurements from a target clinical population with connectome-based WBM promises to become a powerful combination for studying how neuromodulation reconfigures neural circuits to alleviate depressive symptoms. This approach has allowed us to combine, in a unified picture, conventional ERP component-focused TMS-EEG analyses, anti-correlated frontolimbic brain networks, iTBS plasticity-modulated changes to excitation/inhibition balance, and dynamical systems perspectives on how this modulation alters the brain’s attractor landscape.

### Spectral markers of iTBS efficacy

Our spectral analysis on empirical data suggests that iTBS-induced inhibition plays a key role in modulating TMS-induced low-frequency power, as demonstrated by the greater reduction in average 3–10 Hz spectral power in responders compared to non-responders, highlighting this enhanced suppression of theta/alpha activity as the key neurophysiological signature of successful iTBS intervention. Interestingly, this observed effect on empirical data was related to specific changes in synaptic connectivity within our model. The reduction in pyramidal drive to excitatory cells represents a mechanism that shifts the overall E/I balance toward a state of enhanced inhibitory control. This integrated perspective helps reconcile our findings with both the broader literature on GABA-mediated inhibition in depression and the specific synaptic changes observed in our computational model. Indeed, previous research has shown that increased inhibitory control can reduce low-frequency oscillations, a response also noted in animal studies using optogenetic and pharmacological interventions to enhance inhibitory activity^46,47^. In clinical TMS contexts, lower low-frequency power has been linked to treatment efficacy across subjects^48,49^, suggesting that enhanced inhibition may signal a positive response to iTBS. In addition, our simulations demonstrate that increasing inhibition (by reducing P→E connectivity) leads to a marked reduction in low-frequency power, aligning with empirical data and suggesting a mechanistic link between heightened inhibition and observed spectral reductions in severe MDD cases. This finding is consistent with previous modeling studies showing that oriens-lacunosum moleculare (O-LM) cells^50^, key inhibitory interneurons in the hippocampus, adjust their inhibitory response to different frequencies depending on channel types: cells with hyperpolarization-activated cyclic nucleotide-gated (HCN) channels, which play essential roles in regulating cell excitability, are more responsive at higher theta frequencies (4–9 Hz), whereas cells lacking these channels respond more at lower theta (2–5 Hz)^51^. Computational models further support these findings, demonstrating that increased inhibition can reduce low-frequency spectral power, mimicking the patterns observed in our responders^52^.

### Enhanced inhibitory plasticity tracks clinical improvement

In addition to empirical data features, identifying specific biomarkers predicting the likelihood of higher or lower responsiveness to a given treatment or therapy is a key goal in WBM of non-invasive brain stimulation^29,53,54^. We have found, in a data-driven fashion, a significant negative linear relationship between the inhibitory feedback loop before the iTBS protocol and the clinical scores after the treatments for responders. Interestingly, when examining the pre-versus post-iTBS distribution of the inhibitory feedback loop, it was observed that the parameters increased in responders post-treatment and decreased in non-responders. Consistent with this, we also observed a post-iTBS treatment increase in inhibitory activity for responders only, as indicated by the shift in their E/I balance. These findings are consistent with previous modeling work on TBS, which highlights the critical role of pulse count in determining excitatory or inhibitory outcomes. According to the calcium-dependent plasticity (CaDP) model^55^, synaptic changes are governed by intracellular calcium dynamics, with different calcium levels leading to either long-term potentiation (LTP) or long-term depression (LTD). Under this model, the standard 600-pulse iTBS protocol typically produces canonical increases in excitability. However, for pulse counts lower or higher than this— such as 300 or 1200 pulses—the model predicts a reversal of these effects, favoring inhibitory outcomes instead^56^ - consistent with prior experimental results on motor system plasticity effects with variable protocol lengths^57^. In our trial, participants received 1200 pulses per day and, interestingly, despite the high total pulse count, we observed post-treatment inhibition in responders. This suggests that inhibitory outcomes may still emerge under high-dose protocols, potentially due to individual differences in calcium dynamics, cortical state, or metaplasticity mechanisms. These factors, including the temporal spacing of stimulation, could modulate synaptic plasticity in complex and nonlinear ways. Altogether, this suggests that individual differences in the direction of synaptic change are influenced by both the pulse count and individual physiology, such as the phase of calcium oscillations during stimulation, cortical structure, or genetic factors affecting neuronal excitability.

Taken together, these results suggest that the observed alterations in inhibitory synaptic weight, coupled with the shift in E/I balance following successful iTBS treatment, reflect a potential mechanism underlying the therapeutic efficacy of iTBS in depression. Specifically, the increase in inhibitory activity post-treatment in responders may contribute to the restoration of the disrupted E/I balance associated with MDD pathology. Consistent with this, lower levels of mGluR5 (metabotropic glutamate receptor 5)^58^ expression in the prefrontal cortex, particularly in Brodmann’s area 10, have been observed in depressed patients^59^, potentially contributing to disrupted E/I balance in these regions. This highlights the importance of considering the neurophysiological mechanisms involved in E/I balance regulation when understanding and predicting individual responses to neuromodulatory interventions like iTBS. Furthermore, these findings underscore the promise of integrating computational modeling with empirical data to unravel the complex dynamics of brain stimulation therapies. The data-driven discovery in the present study of a relationship between pre-iTBS inhibitory feedback loops and clinical outcomes, coupled with post-treatment increases in inhibitory activity in responders, suggests that inhibitory synaptic changes may play a decisive role in determining treatment success. This insight could inform the development of personalized treatment strategies for depression and related neuropsychiatric conditions, where specific biomarkers, such as E/I balance, guide individualized interventions. It is important to highlight that the number of pulses required for changes in cortical excitability following iTBS may vary between individuals, depending on how TMS interacts with excitatory cortical populations and metaplastic mechanisms. Structural differences in brain convolutions or genetic variations in physiology may cause the same stimulation protocol to activate different groups of neurons across individuals. This variability, combined with the observation from prior experimental^57^ and modeling^56^ work and consistent with our own results that 1200 pulses lead to inhibition, aligns with the idea that adjusting the number of pulses or TMS intensity could minimize differences in treatment response. By identifying and understanding the physiological elements that most influence TMS response, we may be able to develop personalized models that optimize stimulation parameters. These models would allow for the tailoring of clinical treatments to an individual’s neurophysiological profile, ultimately enhancing therapeutic outcomes for depression and other neuropsychiatric disorders. In this way, combining computational models with empirical data offers a powerful approach for refining neuromodulation techniques and improving the precision of brain stimulation therapies.

### Emergence of therapeutically optimal brain states

Furthermore, one of the key advantages of physiologically-based brain modelling is the potential for making meaningful connections between major empirical data features and the physiological constructs instantiated in the model’s states. Our models revealed that a specific brain state, expressed as an eigenvector loading over brain regions and a corresponding scalp topography at the EEG channel level (Fig. 3A), was predictive of the difference in clinical outcomes between responders and non-responders. Notably, the left DLPFC and left sgACC in this brain state exhibited eigenvector loadings of opposite sign. The sgACC is a region positioned at the anterior-inferior end of the cingulum bundle, with extensive connections across prefrontal and limbic structures that have been implicated in depression^60^, and has been linked to clinical response across a diverse range of antidepressant treatment modalities^61–63^. Recent studies have shown that antidepressant outcomes were better when stimulation was delivered at sites of the DLPFC that displayed stronger negative (anticorrelated) FC with the sgACC^24^, a finding that has been replicated across 3 geographically distinct clinical cohorts, different populations, methodologies, scanners, stimulators, and DLPFC targeting approaches^22–25^. Moreover, a 60% to 70% reduction in depressive symptoms occurred when individuals were stimulated near the DLPFC site of maximal FC anticorrelation with sgACC, while those stimulated farther away showed no response or worsening of depressive symptoms^23^. The fact that similar topographic maps to these fMRI sgACC anticorrelation patterns were obtained from our brain network model of TMS-EEG responses in MDD patients is an intriguing and unanticipated result. Our temporal analysis of sgACC activation dynamics (Fig. 3C) showed significantly higher sgACC engagement patterns in responders compared to non-responders at baseline, prior to iTBS intervention. This finding aligns with the therapeutic mechanism of iTBS, which aims to modulate sgACC activity through stimulation of the DLPFC^64,65^. This suggests that patients with more robust sgACC engagement pre-iTBS may possess the necessary neurophysiological substrate for effective modulation via DLPFC stimulation. This observation supports the network-based conceptualization of iTBS efficacy, where the primary target (DLPFC) serves as an entry point to influence deeper limbic structures, particularly the sgACC, through existing functional connections. Moreover, these findings extend our understanding of the DLPFC-sgACC relationship beyond static connectivity patterns to include the dynamic temporal characteristics of sgACC recruitment. The significantly higher normalized sgACC engagement in responders suggests that pre-treatment sgACC excitability may be a crucial determinant of iTBS efficacy. This could explain why stimulation of DLPFC regions with stronger negative functional connectivity to sgACC yields better clinical outcomes^23,24,45^ – these connections may facilitate more effective modulation of an already sensitized sgACC in treatment-responsive individuals.

Our neural trajectory analysis (Fig. 4) provides compelling evidence that successful iTBS treatment alters the dynamics of cortical circuits in a way that distinguishes responders from non-responders. The three-dimensional visualization of neural trajectories reveals that while both groups initially exhibit similar dynamics pre-treatment, responders show a marked trajectory contraction post-iTBS, suggesting increased stability in their attractor dynamics (Fig. 4A). This stabilization is quantified by the significantly greater trajectory shifts in responders (Fig. 4B) and their consistently closer adherence to the fixed point attractor following treatment (Fig. 4C).

These findings suggest that iTBS may exert its therapeutic effects by reconfiguring the dynamical landscape of prefrontal-limbic circuits, potentially restoring a more stable, less chaotic pattern of neural activity. The greater magnitude of trajectory reorganization in responders indicates that a substantial shift in neural dynamics may be necessary for clinical improvement. Simultaneously, the closer proximity to the attractor point post-treatment suggests that effective iTBS therapy creates a more stable neural state that is less susceptible to perturbations—a characteristic that may contribute to symptom relief by dampening the excessive reactivity often observed in depressive states^66,67^.

This dynamical systems perspective complements our findings regarding E/I balance and sgACC-DLPFC interactions, providing a more comprehensive framework for understanding iTBS efficacy. The ability of iTBS to induce meaningful reconfiguration of attractor dynamics appears to be a critical determinant of treatment response, potentially serving as another neurophysiological signature that could guide personalized neuromodulation approaches.

### Limitations and future directions

Despite our promising findings, several limitations should be acknowledged. First, our computational model, while detailed, necessarily represents a simplification of the complex neural dynamics underlying TMS responses. The Jansen-Rit neural mass formulation captures population-level activity but cannot resolve cellular-level mechanisms, or mechanisms extending beyond the minimal JR three-population circuit that may be relevant to iTBS effects. Second, our structural connectivity data was derived from a normative dataset rather than subject-specific tractography, which may not fully capture individual variations in anatomical connections relevant to treatment response. Third, the 1200-pulse iTBS protocol used in this study differs from the standard 600-pulse protocol commonly used in clinical settings, potentially limiting direct comparison with other clinical studies. Fourth, while our approach identified significant associations between model parameters and clinical outcomes, the causal relationship between these parameters and therapeutic effects requires further experimental validation. Fifth, the absence of a sham control condition in our study design limits our ability to distinguish between specific neurophysiological effects of iTBS and non-specific factors such as placebo effects or natural disease fluctuations. Sixth, our analyses focused primarily on prefrontal-subgenual interactions and cortical E/I balance, but other neural circuits and mechanisms not captured in our model may also contribute to treatment outcomes. Finally, although our sample size was substantial for a TMS-EEG study, larger and more diverse patient cohorts would be valuable to validate the generalizability of our findings across different TRD populations. Future studies combining longitudinal TMS-EEG with individualized connectivity measures, sham controls, and more sophisticated computational models could address these limitations and further refine our understanding of the neurophysiological mechanisms underlying iTBS efficacy In conclusion: our results, and the framework for investigating the scientific questions we are introducing here, not only enhance our understanding of the therapeutic mechanisms of iTBS, but also highlight the potential for developing neurophysiologically-informed biomarkers to guide personalized neuromodulation treatments. As computational psychiatry continues to evolve, the integration of biophysically-based modeling with multimodal neuroimaging promises to transform our approach to treatment-resistant depression, moving toward precision interventions tailored to individual neural dynamics. Future work building on these findings could lead to optimized stimulation protocols that maximize clinical benefits by targeting specific neurophysiological mechanisms, ultimately improving outcomes for patients with this debilitating condition.

## CONCLUSION

Our study demonstrates that successful iTBS treatment for depression is characterized by two key neurophysiological mechanisms: enhanced inhibitory control (evidenced by greater reductions in low-frequency oscillatory power and increased I→P connectivity) and specific pre-treatment frontolimbic connectivity patterns between DLPFC and sgACC. These findings advance our understanding of how neuromodulation alters brain circuit dynamics in treatment-resistant depression, suggesting potential predictive biomarkers and opening avenues for personalized TMS interventions.

## METHODS

The analyses conducted in the present study consist of four main components: (i) EEG preprocessing and calculation of TMS stimulation-evoked responses, (ii) construction of anatomical connectivity priors for our computational model using diffusion-weighted MRI tractography, (iii) simulation of whole-brain dynamics and stimulation-evoked responses with a connectome-based neural mass model, and (iv) fitting of the model to individual-subject scalp EEG data, statistical comparisons of the resultant estimated physiological parameters alongside other clinical variables, and further analysis of simulated activity properties. A schematic overview of the overall approach is given in Fig. 5.

**Fig. 5.**
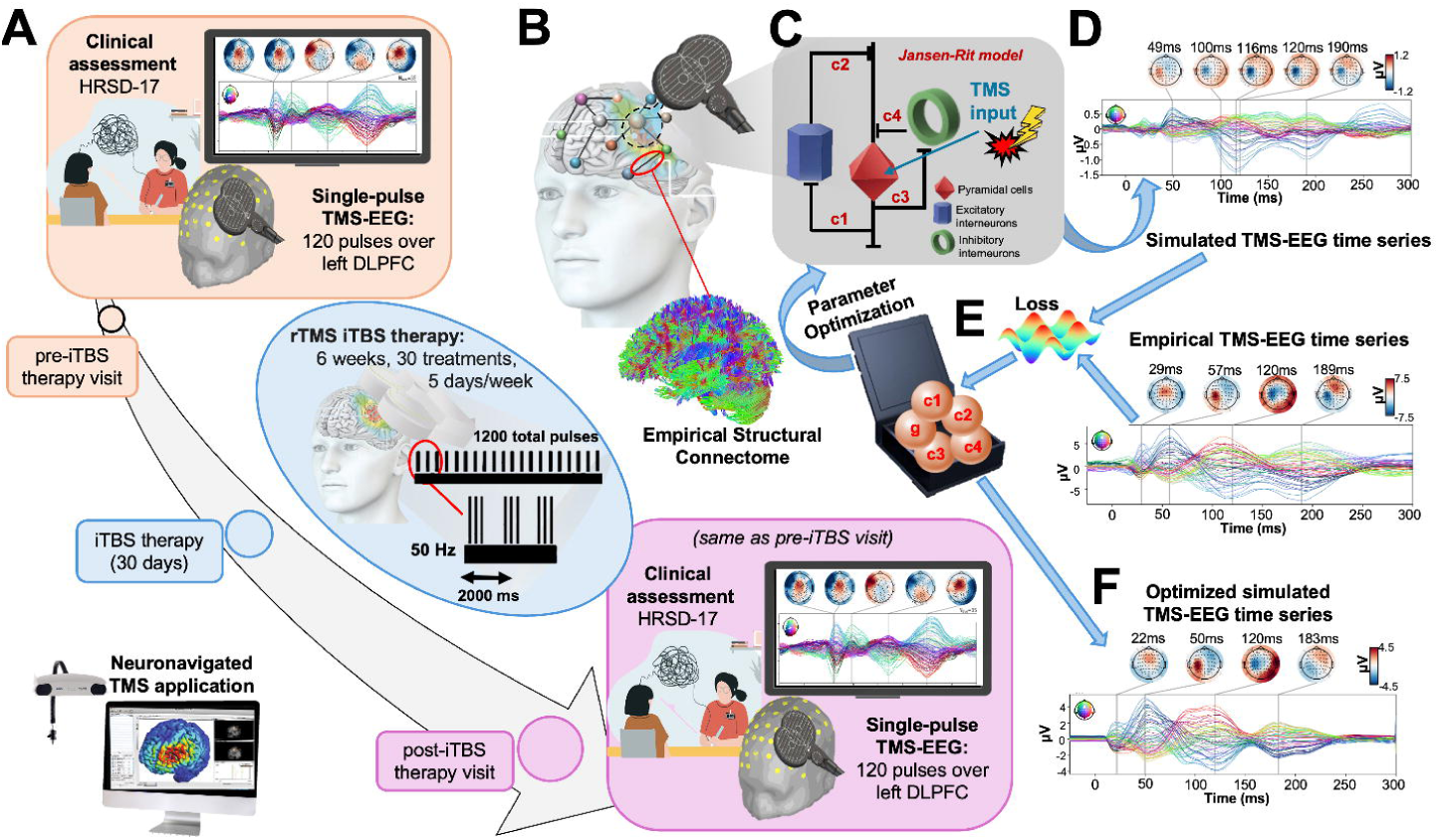
Overview of study design and methodological workflow for subject-specific connectome-based whole-brain modeling of TMS evoked potentials. **(A)** Single-pulse TMS-EEG evoked responses and HRSD-17 depression scale were recorded before and after 30 days’ rTMS iTBS therapy. The iTBS protocol consisted of daily treatment for 6 weeks (30 sessions, 5 days/week) applied over the L-DLPFC, with 1200 total pulses delivered per day. For further details on the data collection and EEG preprocessing methodology see Strafella et al. (2023)^42^. **(B)** Diffusion-weighted MRI tractography was computed from a sample of healthy young individuals from the HCP Dataset^68^, and then averaged to give a grand-mean anatomical connectome. **(C)** The Jansen-Rit model^38^ was embedded was embedded in each of the 200 nodes of the Schaefer atlas^69^ for simulating and fitting neural activity time series. The TMS-induced depolarization of the resting membrane potential was modeled by a perturbing voltage offset to the mean membrane potential of the pyramidal cell. **(D)** A lead field matrix was then used for moving the parcels’ time series into channel space and generating simulated TMS-EEG. **(E)** The goodness-of-fit (loss) was calculated between simulated and empirical TMS-EEG time series. (E) Utilizing the autodiff-computed gradient^70^ between the objective function and model parameters, model parameters were optimized using the ADAM algorithm^71^. **(F)** Finally, the optimized model parameters were used to generate the fitted, simulated TMS-EEG activity, for which we report comparisons with the empirical data at both the channel and source level using conventional statistical techniques.

### Recruitment and trial design

As part of a triple-blind randomized controlled trial conducted at the Centre for Addiction and Mental Health, University Health Network, and the University of British Columbia, 90 participants from these sites underwent TMS-EEG assessments both at baseline and after an iTBS treatment (clinicaltrials.gov identifier NCT02729792)^72^. Eligibility criteria included being between the ages of 18 and 59, meeting the MDD diagnosis criteria based on the Mini-International Neuropsychiatric Interview, having a baseline HRSD-17 score of 18 or higher (indicating moderate-to-severe depression), maintaining stable psychotropic medications for at least 4 weeks prior to screening, and either failing to respond to one adequate antidepressant trial or being unable to tolerate two different antidepressant trials. This recruitment approach specifically targeted patients with treatment resistance, a population with significant unmet clinical need. The study was approved in accordance with the Declaration of Helsinki, and all participants provided written informed consent. For details regarding the patient recruitment, the data acquisition and the preprocessing steps, and the iTBS protocol, please refer to our paper^42^ and supplementary materials. All the preprocessed EEG analyses in the present work were performed using the MNE software library^73^ (mne.tools/stable/index.html) running in Python 3.6.

### Overview of computational modelling approach

In our study, we employed a WBM approach to analyze pre- and post-iTBS EEG data from a cohort of 90 patients with MDD. This model incorporated 200 distinct cortical regions based on the Schaefer 200-parcel, 7-network atlas^69^, connected with a set of inter-regional weights derived from the anatomical connectome. These connectivity weights were obtained by averaging diffusion MRI tractography data from 400 subjects in the Human Connectome Project (HCP) dataset^68^. Jansen-Rit (JR) neural mass dynamics^38^ at each modeled region described the process of stimulated activation and damped oscillatory responses resulting from local interactions within cortical microcircuits, with these effects propagating to regions distal to the stimulated site via the anatomical connectome. After specifying its structure and a common set of priors for all parameters, the model was fit to EEG data separately for each patient. This resulted in a set of individualized physiological and connectivity parameters, having a mechanistic causal influence on several spatial and temporal features of the brain stimulation response, which we subsequently interrogated to obtain further insight into our research questions around possible differences between patients who reported benefits from iTBS and those who did not. For details regarding the computational model and the estimation of parameters, please refer to Momi et al.^35,36^ and supplementary material and methods. For a graphical overview of all optimized parameter distributions, please refer to Supplementary Fig. S1.

### Assessing the similarity between simulated and empirical evoked responses

To further assess the goodness-of-fit the simulated waveforms arrived at after convergence of the ADAM algorithm^71^, Pearson correlation coefficients and corresponding p-values between empirical and model-generated waveforms were computed for each subject. To control for type I error, this result was compared with a null distribution constructed from 1,000 time-wise random permutations, with a significance threshold set at p<0.05. For an overview of the goodness-of-fit between simulated and empirical TMS-EEG data, including representative butterfly plots and correlation distributions across subjects, please refer to Supplementary Fig. S2.

### Stimulus-induced spectral power analyses

For each subject’s pre-iTBS and post-iTBS recordings, we computed stimulus-induced spectral power across frequencies from 2 Hz to 50 Hz, analyzing differences attributable to iTBS treatment. A Morlet wavelet was created for each frequency of interest and convolved with hd-EEG data for each channel. We then calculated the power spectrum, applied a logarithmic transformation, and averaged these power values across trials within the defined analysis window (0–300 ms), selected to capture the primary induced response. For relative power, values were normalized to baseline power calculated over a baseline window of -300 to 0 ms.

Post-iTBS versus pre-iTBS comparisons were performed for each subject using condition-wise permutation testing and cluster-based thresholding for multiple comparisons. Specifically, the permutation test converted the difference between induced and baseline windows into z-scores, based on a null distribution generated by 1,000 permutations with random window label swaps. These z-scores were thresholded at p < 0.05. An additional 1,000 iterations of the permutation test yielded a distribution of cluster sizes under the null hypothesis, identifying time-frequency clusters exceeding the 95th percentile for significance. Finally, we obtained the subject- and session-specific stimulus-induced spectral power by averaging across channels. For an overview of the group-level TMS-induced spectral power changes following iTBS treatment for responders and non-responders, please refer to Supplementary Fig. S3. To compare responders and non-responders, we performed a second-level statistical analysis. The significant z-score maps from the first-level analysis were grouped by treatment outcome, and differences between responders and non-responders were assessed using another permutation test (1,000 iterations) with cluster-based correction (Fig. 1A). This hierarchical statistical approach allowed us to identify spectro-temporal patterns that significantly differed between treatment outcome groups while controlling for multiple comparisons at both the individual and group levels.

A key advantage of physiologically-based brain modeling is its ability to link key empirical data features with physiological constructs represented by the model’s parameters. In this study, we leveraged this approach by investigating the relationship between the significant clusters in the stimulus-induced time-frequency spectral power maps and the physiological parameters of the Jansen-Rit model (Fig. 1B). To conduct this analysis, we focused on significant clusters within the stimulus-induced power spectra, which represents frequency ranges where iTBS-induced changes were most prominent. For each subject, we extracted the average spectral power values from this significant region and computed the difference between post-iTBS and pre-iTBS values. This subtraction (post-iTBS minus pre-iTBS) allowed us to quantify the degree of iTBS-induced modulation within the significant spectral region. This approach enabled us to explore direct associations between spectral power changes and underlying physiological mechanisms modeled in the Jansen-Rit framework (Fig. 1C, 1D & 1E), offering insights into the neural dynamics influenced by iTBS.

### Evaluation of iTBS-induced changes in physiological model parameters and E/I balance

We investigated the effects of iTBS treatment on several key JR model parameters associated with synaptic dynamics and connectivity (Fig. 2A & 2B). The parameters analyzed were the synaptic time constant of the excitatory population (a), the synaptic time constant of the inhibitory population (b), and the synaptic weights for pyramidal-to-excitatory (c1), excitatory-to-pyramidal (c2), pyramidal-to-inhibitory (c3) and inhibitory-to-pyramidal (c4) populations. The c1-c4 labels are the standard notation used for JR model, however as more intuitive shorthand we also use the substitutions P→E, E→P, P→I, and I→P for c1, c2, c3, and c4, respectively. As an additional summary of the physiological and dynamical state of each subject’s pre- and post-iTBS brain activity, we also explored excitatory-inhibitory (E/I) balance (Fig. 2C), defined as [c1*c2] / [c3*c4] (i.e. the ratio of the combined gains of the JR excitatory and inhibitory feedback loops), where the c parameters were first mean-normalized across all participants to ensure comparability across conditions. Specifically, for each parameter (c1-c4), we calculated the mean value across all participants and all conditions (pre-iTBS and post-iTBS), then divided each individual subject’s parameter values by this global mean. This normalization ensures that all parameters contribute proportionally to the E/I ratio based on their relative values rather than being dominated by parameters with larger absolute magnitudes. Statistical analyses on estimated model parameters were performed using R-Studio Version 2024.04.2+764. For each of the JR parameters and the composite E/I balance metric, we conducted a series of 2x2 repeated measures ANOVAs with “time” (two levels: pre-iTBS and post-iTBS) as a within-subject factor and “group” (two levels: responders and non-responders) as a between-subject factor. This approach allowed us to assess how treatment and time interacted to affect the specified physiological parameters. Additionally, post-hoc paired t-tests were employed to detect changes in parameter values from pre-iTBS to post-iTBS for both responders and non-responders.

### Predicting iTBS clinical outcome using model-derived Hidden Brain States

We employed a systematic approach to investigate the relationship between model-derived brain states and clinical response to iTBS. First, we extracted the time series of TMS-evoked activity (-100-300ms) from the optimized JR model for each of the three neuronal populations (pyramidal cells, excitatory interneurons, and inhibitory interneurons), for all 200 cortical regions. For each subject, we then concatenated these three arrays in time and performed dimensionality reduction using singular value decomposition (SVD), obtaining principal components representing dominant spatiotemporal patterns of neural activity.

Using these pre-treatment components, we applied mean-centered PLS analysis^74,75^ to identify brain states that maximally discriminate between responders and non-responders (Fig. 3A). This multivariate statistical approach identifies latent variables (brain states) that maximize the covariance between neural activity patterns and group membership, while accounting for within-group variability. Statistical significance was assessed through permutation testing (5,000 permutations), randomly reassigning subjects to groups to establish a null distribution. The reliability of regional contributions to the discriminative brain states was evaluated using bootstrap resampling (1,000 samples).

To visualize how the identified brain states differentiated between responders and non-responders, we specifically examined the relationship between loadings in the DLPFC (stimulation target) and sgACC (key region implicated in depression pathophysiology) for each subject (Fig. 3B). This approach allowed us to quantify the expression of the treatment-predictive brain state pattern at the individual subject level and identify neurophysiological signatures that may serve as biomarkers for treatment response.

For temporal dynamics analysis, we extracted normalized activation time courses of the sgACC from the model for each subject (Fig. 3C). Again, these time courses represented the model-predicted neural response to stimulation, measured from -100 ms pre-stimulation to 300 ms post-stimulation. We calculated group-averaged time courses for responders and non-responders separately, with shaded regions representing standard error of the mean. Statistical comparison between groups was performed using cluster-based permutation testing to identify time windows with significant differences while controlling for multiple comparisons.

To quantify the overall sgACC engagement differences between groups, we calculated the AUC of normalized sgACC activation for each subject (Fig. 3D). AUC values were normalized to account for individual differences in baseline activity. Group differences in AUC values were assessed using independent samples t-tests, with significance set at p < 0.05. Violin plots were used to visualize the distribution of AUC values in each group, with individual data points overlaid to show subject-level variability.

### Neural trajectory analysis in model-derived state space

To visualize and quantify changes in brain dynamics induced by iTBS, we analyzed the neural trajectories in the three-dimensional state space of the JR neural mass model (Fig. 4A). For each group (responders and non-responders) and session (pre- and post-iTBS), we constructed state-space trajectories using the activity of the three neural populations in the model: pyramidal neurons, excitatory interneurons, and inhibitory interneurons. This approach allowed us to examine how the neural dynamics evolve over time in response to stimulation and how these dynamics are altered by iTBS treatment. The reconstructed trajectories were normalized using the group-averaged singular vectors and singular values. To compare dynamics across conditions, we plotted the reconstructed trajectories in 3D space and quantified their geometric properties using two complementary metrics. First, we computed point-wise Euclidean distances between post-iTBS and pre-iTBS trajectories within each group (Fig. 4B), capturing the degree of treatment-induced deviation in neural state space. Second, we computed point-wise Euclidean distances from the fixed point attractor for each post-iTBS trajectory (Fig. 4C). To calculate this fixed point, we numerically identified where the model’s differential equations equal zero using each subject’s fitted parameters, testing a large number of potential values and computing the Jacobian matrix to confirm stability. These distances reflect the stability of the dynamical system, where smaller distances indicate a tighter convergence toward the attractor and reduced sensitivity to perturbations (e.g., TMS pulses).

Statistical differences between responders and non-responders were assessed by comparing the average Euclidean distance curves using a two-sample non-parametric permutation test (1,000 iterations), performed separately for Fig. 4B and 4C. Group-level time-series distances were summarized by their AUC, and the null distribution was generated by randomly permuting group labels to test the hypothesis that iTBS alters attractor geometry more strongly in responders than in non-responders.

## ACKNOWLEDGEMENTS

This work was supported by the CAMH Discovery Fund Talent Competition 2023-24 (D.M.), Krembil Foundation (JG), Canadian Institute of Health Research (JG), Labatt Family Foundation (JG). The CARTBIND study was funded by Brain Canada with the financial assistance of Health Canada, the Temerty Centre for Therapeutic Brain Intervention, Philantropic donation to the NINET lab, Seedlings Foundation, the Krembil Family Foundation, the Arrell Family Foundation, and the Toronto General and Western Hospital Foundation.

