## Supplementary material for "Excitation-Inhibition Balance and Fronto-Limbic Connectivity Drive TMS Treatment Outcomes in Refractory Depression": SI

**1. SUPPLEMENTARY MATERIAL AND METHODS**

1.1 Patient Sample and Exclusion Criteria

1.2 Neuroimaging data and definition of connectome weight priors

1.3 Large-scale connectome-based neurophysiological brain network model

1.4 Individual-subject Jansen-Rit connectome model parameter estimation from hd-EEG data

**2. SUPPLEMENTARY RESULTS**

2.1 Non-significant Effects of iTBS on Physiological Parameters

2.2 Connectome-based neurophysiological modelling accurately reproduces subject-specific TMS-EEG dynamics

2.3 TMS-Evoked Spectral Power in Responders and Non-responders

**3. SUPPLEMENTARY VIDEOS**

Video V1. Dynamic visualization of TMS-EEG neural trajectories pre-iTBS

Video V2. Dynamic visualization of TMS-EEG neural trajectories post-iTBS

**4. SUPPLEMENTARY REFERENCES**

### 1. SUPPLEMENTARY MATERIAL AND METHODS

#### 1.1 Patient Sample and Exclusion Criteria

This randomized controlled trial was conducted at three Canadian academic centers (Centre for Addiction and Mental Health, University Health Network, both affiliated with the University of Toronto, and the University of British Columbia Hospital) with ethics board approval and participant written informed consent. The trial was registered on ClinicalTrials.gov (NCT02729792)[1]. Between April 2016 and February 2018, adults aged 18–59 with a confirmed diagnosis of major depressive disorder (MDD), single or recurrent, were enrolled. Inclusion criteria included failure to respond to at least one adequate antidepressant trial (ATHF score ≥3) or intolerance to two different trials, HRSD-17 score ≥18, stable medication for four weeks prior to enrollment, and eligibility for rTMS based on the TASS questionnaire and normal thyroid function. Key exclusions were recent substance use disorder, unstable medical or neurological conditions, active suicidality, pregnancy, primary comorbid psychiatric disorders (e.g., bipolar, psychotic, anxiety, or personality disorders), prior ECT failure in the current or previous episode, prior rTMS exposure, metal implants near the head, or unstable psychotherapy. Patients using anticonvulsants or high doses of benzodiazepines were also excluded.

Stimulation targeted the left DLPFC using neuronavigation based on the MNI coordinate [–38, 44, 26] during the first session; subsequent sessions used marked scalp locations. Active stimulation was delivered at 120% RMT using a MagPro X100/R30 stimulator with a B70 coil; sham stimulation was delivered at the Pz site using a matched placebo coil. All participants completed 60 sessions across 30 treatment days, with active and sham stimulation matched across groups in dose and timing. Clinical assessments occurred at baseline, every two treatment days for the first 10 sessions, every five sessions thereafter, and at 1, 4, and 12 weeks post-treatment. Adverse events and procedural pain (rated 0–10) were recorded at each session. The primary outcome was change in HRSD-17 from baseline to day 10; secondary outcomes included HRSD-17 change at day 30 and changes in BDI-II and QIDS-SR16 scores. Response was defined as ≥50% reduction in HRSD-17; remission as HRSD-17 <8. Additional biomarkers collected during the trial will be reported separately.

##### 1.2 Neuroimaging data and definition of connectome weight priors

To establish anatomical connectivity priors representative of the population, we conducted DW-MRI tractography reconstructions on a large sample of healthy young individuals (see also [2]). This dataset comprised structural neuroimaging data from 400 healthy young subjects (170 males, aged 21–35 years), sourced from the Human Connectome Project (HCP) Dataset (available at humanconnectome.org/study/hcp-young-adult)[3].

Our DW-MRI preprocessing workflow was executed on Ubuntu 18.04 LTS using tools from the FMRIB Software Library (FSL 5.0.3; https://www.fmrib.ox.ac.uk/fsl)[4], MRtrix3 (https://www.MRtrix.readthedocs.io)[5], and FreeSurfer 6.0[6]. These images were already corrected for motion using FSL's EDDY[7] as part of the HCP minimally-preprocessed diffusion pipeline[8]. We estimated the multi-shell multi-tissue response function using constrained spherical deconvolution[9] and segmented T1-weighted (T1w) images, which were already coregistered to the b0 volume, using the FAST algorithm[10]. We employed anatomically-constrained tractography to generate an initial tractogram with 10 million streamlines using second-order integration over fiber orientation distributions[11]. Subsequently, we applied the spherical-deconvolution informed filtering of tractograms (SIFT2) methodology to yield more biologically accurate measures of fiber connectivity[12].

To define brain regions or network nodes, we utilized the 200-region atlas developed by Schaefer et al. This atlas was mapped to each individual's FreeSurfer surfaces using spherical registration[13]. Additionally, this atlas provided categorical assignments of regions into seven canonical functional brain networks (Visual network: VN, Somatomotor network: SMN, Dorsal attention network: DAN, Anterior salience network: SN, Limbic network: LN, Fronto-parietal network: FPN, Default mode network: DMN).

By combining this atlas with the filtered streamlines, we derived 200 × 200 anatomical connectivity matrices. These matrices contained information on the number of streamlines and fiber length connecting each pair of regions. We then averaged these connectomes across the 400 HCP subjects, resulting in a population-representative connectome matrix.

To prepare this matrix for physiological network modeling, we rescaled the values. This rescaling involved two steps: first, taking the matrix Laplacian, which set each row sum (i.e., each node's weighted in-degree) to zero by subtracting row sums from the diagonal; and second, performing scalar division of all entries by the matrix norm. This ensured the linear stability of the matrix, with all eigenvalues having a negative real part except for one eigenvalue, which was zero. The Laplacian approach has been commonly employed in previous research on whole-brain modeling[14–16].

##### 1.3 Large-scale connectome-based neurophysiological brain network model

The methods for modeling whole-brain dynamics are highly similar to those in Momi et al.[2,17]. Our brain network model encompasses 200 cortical areas, each representing the population-averaged activity of an individual brain region in line with mean-field theory principles[18]. In describing the activity at each node, we employed the Jansen-Rit (JR) equations, a widely adopted neurophysiological model applied to both stimulus-evoked and resting-state EEG activity measurements[19–21], that we have also recently employed for modelling evoked responses to TMS[2] and intracerebral electrical stimulation[17]. The network model connectivity incorporated pyramidal-to-inhibitory, pyramidal-to-excitatory, and inhibitory-to-excitatory interneuron connections[19].

The JR model serves as a relatively coarse-grained representation of the cortical microcircuit, consisting of three interconnected neural populations: pyramidal projection neurons, excitatory interneurons, and inhibitory interneurons. While the excitatory and inhibitory populations both receive input from, and provide feedback to the pyramidal population, they do not interact with each other directly. Consequently, the circuit motif comprises one positive and one negative feedback loop. Within each of these three neural populations, the post-synaptic somatic and dendritic membrane response to incoming action potentials is described by second-order differential equations.

$$\ddot{v}(t) + \frac{2}{\tau_{e,i}}\dot{v}(t) + \frac{1}{\tau_{e,i}^2}v(t) = \frac{H_{e,i}}{\tau_{e,i}}m(t) \quad (1)$$

which is equivalent to a convolution of incoming activity with a synaptic impulse response function

$$v(t) = \int_0^\infty d\tau m(\tau) \cdot h_{e,i}(t-\tau) \quad (2)$$

whose kernel $h_{e,i}$ is given by

$$h_{e,i} = \frac{H_{e,i}}{\tau_{e,i}} \cdot t \cdot exp\left(-\frac{t}{\tau_{e,i}}\right) \quad (3)$$

where $m$ is the (population-average) presynaptic input, $v$ is the postsynaptic membrane potential, $H_{e,i}$ is the maximum postsynaptic potential, and $\tau_{e,i}$ a lumped representation of delays occurring during the synaptic transmission.

The synaptic response function, often referred to as a pulse-to-wave operator following Freeman's terminology[22], plays a crucial role in regulating the excitability of the neural population. Of specific relevance to our current study are the time constants $\tau_e$ and $\tau_i$, which govern this excitability.

In addition to the pulse-to-wave operator for synaptic responses, each neural population incorporates a wave-to-pulse operator that determines the population's output, represented by the (population-average) instantaneous firing rate. The firing rate is a function of the somatic membrane potential and follows a sigmoidal pattern,

$$S(v) = \frac{e_0}{1 - exp\left(r(v_0 - v)\right)} \quad (4)$$

where $e_0$ is the maximum firing rate, $r$ is the steepness of the sigmoid function, and $v_0$ is the postsynaptic potential for which half of the maximum firing rate is achieved.

In practice, as is standard with the JR model, we express the three sets of second-order differential equations that follow the structure of Equation 1 as three sets of coupled first-order differential equations. As a result, the complete JR system for each individual cortical area, denoted as j ∈ {i, i+1,..., N} within our network comprising N=200 regions, can be represented by the following six equations:

$$\dot{v}_{j1} = x_{j1} \quad (5)$$

$$\dot{x}_{j1} = \frac{H_e}{\tau_e}\left(C_1 S\left(C_2 v_{j3}\right) + g_f \sum_{i=0}^{N} wf_{ji}\left(v_{i3}(\mathrm{t} - m_{ji}) - v_{j3}\right)\right) - \frac{2}{\tau_e} x_{j1} - \frac{1}{\tau_e^2} v_{j1} \quad (6)$$

$$\dot{v}_{j2} = x_{j2} \quad (7)$$

$$\dot{x}_{j2} = \frac{H_i}{\tau_i}\left(C_4 S\left(C_3 v_{3j}\right) + g_b \sum_{i=0}^{N} wb_{ji}\left(v_{i3}(\mathrm{t} - m_{ji}) - v_{j3}\right)\right) - \frac{2}{\tau_i} x_{j2} - \frac{1}{\tau_i^2} v_{j2} \quad (8)$$

$$\dot{v}_{j3} = x_{j3} \quad (9)$$

$$x_{j3} = \frac{H_e}{\tau_e}\left(S\left(v_{j1} - v_{j2}\right) + g \sum_{i=0}^{N} wl_{ji}\left(v_{i3}(\mathrm{t} - m_{ji}) - v_{j3}\right) + p_j\right) - \frac{2}{\tau_e} x_{j3} - \frac{1}{\tau_e^2} v_{j3} \quad (10)$$

where $v_{1,2,3}$ is the average postsynaptic membrane potential and $x_{1,2,3}$ are the mean currents of the excitatory interneuron, inhibitory interneuron, and pyramidal cell populations, respectively. Here, $g$ denotes the global coupling strength, while $W$ represents the structural connectome, defining the anatomical connectivity between regions $i$ and $j$ across the network.

Importantly, due to the finite velocity of long-range axonal conduction, these inputs exhibit delays ranging from approximately 5 to 50 ms, with variations specific to each connection depending on their physical length. These temporal lags are determined by $m_{ji}$, denoting the $j, i_{th}$ entry in the delays matrix $M$, calculated as $T / s$. Here, $T$ represents the inter-regional fiber tract length matrix, and s denotes the global axonal conduction velocity.

Of particular significance in the present work, we modeled the stimulus-evoked depolarization of the resting membrane potential by introducing an external perturbing voltage offset, denoted as $p_j$, which is applied to the pyramidal interneuron population.

To identify the brain regions engaged by DLPFC-targeted TMS, and therefore which nodes in our simulation to inject an external input, and with what magnitude, the TMS-induced electric field was modeled with SimNIBS[23,24] (simnibs.github.io/simnibs). A tetrahedral head model (mesh file) was created, consisting of five tissue types: white matter (WM), gray matter (GM), cerebro-spinal fluid (CSF), skull, and scalp. The assigned conductivity values were fixed: 0.126 S/m (WM), 0.275 S/m (GM), 1.654 S/m (CSF), 0.01 S/m (skull), 0.465 S/m (scalp). The distance between the coil and the cortex was set to 10 mm using the standard MNI152 brain model in SimNIBS, with the coil handle oriented according to established DLPFC targeting coordinates following the methods and materials of the original paper of [25,26]. The rate of change of the coil current (dI/dt) was calculated assuming a quasistatic regime[26], according to the following equation:

$$E = \frac{\partial A}{\partial t} - \Delta\varphi \quad (11)$$

where E represents the electric field vector, A is the magnetic vector potential of the TMS coil, ∂A/∂t captures the time-dependent changes in the magnetic vector potential, and Δφ denotes the electric potential gradient. The normalized electric field or E-field distribution, modeled with SimNIBS in the MNI152 standard-space, was thresholded at 83% of its maximal value, following recent estimates of the E-field thresholds above which tissue is activated by TMS[27]. This thresholded E-field map was then used to inject a weighted stimulus into the target regions in the model.

The channel-level hd-EEG signals in our model were derived by computing the difference between the excitatory *($v_1(t)$)* and inhibitory *($v_2(t)$)* interneuron post-synaptic potentials (corresponding to the net somatic membrane potential of the pyramidal cell population[28]) at each cortical parcel, and then projecting these signals into the hd-EEG channel space[29].

$$Y = G \cdot X + \epsilon \quad (12)$$

where $Y$ is the channel-level hd-EEG signals and $G$ is the lead field matrix derived by averaging the sensor-level forward solution over cortical regions.

##### 1.4 Individual-subject Jansen-Rit connectome model parameter estimation from hd-EEG data

We employed a recent technique[2,17] for optimizing parameters in our brain network model when fitting individual-subject evoked potential waveforms and extracting subject-specific physiological parameters from empirical data. A notable component of this methodology is our implementation of the connectome-base neural mass model described above in PyTorch[30], a machine learning software library that is widely used in both academic and commercial sectors. In order to systematize our and others' work with this novel approach, we have recently developed a new Python library, Whole-Brain Modelling in PyTorch (WhoBPyT), available at github.com/griffithslab/whobpyt, which implements and documents the basic technique and our applications of it to resting and stimulus/task-evoked fMRI, EEG, MEG, sEEG and fNIRS data.

Transitioning to the PyTorch-based framework from more conventional numerical simulation libraries required some minor adjustments to accommodate tensor data structures and greater memory load. However, this shift offers a significant advantage by seamlessly accommodating gradient-based parameter optimization through automatic differentiation-based algorithms. This is particularly useful for handling complex sets of equations that do not yield easily computable Jacobians. Our approach falls in line with a growing trend[31,32], where the parallels between physiologically-based large-scale brain network models and deep recurrent neural networks in machine learning prove both technically and conceptually valuable.

Recently, we have successfully applied this technique to fast-timescale evoked responses to dissect the spatio-temporal physiological origin of TMS-evoked potentials[2] and iEEG-evoked potentials[17].

The optimization algorithm operates by segmenting a subject's multi-channel evoked potential waveform, which spans 400 ms (from -100 ms to +300 ms post-stimulus) and is trial-averaged, into discrete, non-overlapping windows of 20 ms, referred to as "batches." The entire simulation covers a duration of 400 ms, encompassing a 100 ms baseline period before the electric pulse and a 300 ms period after the introduction of stimulation. Prior to the commencement of the 100 ms baseline, a 20 ms burn-in period is included to allow the system to stabilize following the initial transient caused by randomly assigned initial conditions for the state variables.

Iterating through each batch in the time series sequentially, we generated the JR model-simulated evoked potentials $\hat{y}$ using the current parameter values and forward-Euler integration of equations 5-10. We then assessed its alignment with the empirical evoked potentials y by employing the mean-squared error (MSE) loss function

$$L = \frac{1}{N_t}\sum_{t=1}^{N_t}\left(\frac{1}{N_{ch}}\sum_{i=1}^{N_{ch}}\left(y_i(t) - \hat{y}_i(t)\right)^2\right) \quad (13)$$

where $N_t$ is the number of the time points, $N_{ch}$ is the number of hd-EEG channels, and $L$ is the total loss. It is assumed that the model parameters are Gaussian. Together with a complexity-penalizing regularization term on each model parameter $\theta$,

$$C = ln\,\sigma + \frac{1}{\sigma^2}(\theta - \mu)^2 \quad (14)$$

Here, the mean *($\mu$)* and standard deviation *($\sigma$)* of the model parameter $\theta$ serve as optimization hyperparameters. Equation 14 characterizes the complexity of the model parameters, serving as a regularization term to prevent overfitting and enhance model robustness. The loss function $L$ and the complexity term $C$ are combined into a final objective function,

$$O = L + \lambda C \quad (15)$$

where, λ is a regularization parameter that balances the influence of the complexity term relative to the loss function. This combined objective function is then supplied to PyTorch's native stochastic gradient descent-based algorithm ADAM for optimization[33]. As the batch window traverses the entirety of the evoked potential time series, it cyclically reverts to the beginning and repeats until convergence is achieved. Each repetition of this loop is, following ML convention, termed an *Epoch* (but not to be confused with an Epoch in the EEG trial sense). Upon optimization completion, the average of each parameter's values over the last 100 batches give its final estimated value which we use both for statistical analyses and for further simulations.

We estimated five sets of parameters for each subject: i) physiological parameters defining the dynamic regime of the JR cortical microcircuit, ii) connection weight modulations, iii) lead field modulations, iv) conduction delays, v) hyperparameters. The first of these is the most important as it is the primary determinant of the network nodes' waveform shape and oscillatory characteristics of their stimulation responses, and in the present context pertains to individual differences in baseline neurophysiology. The second allows for individual variation away from the tractography-based population-representative prior values on anatomical connection strengths described above. It is however important here to strike a balance between empirically-motivated constraints and flexibility during fitting. Our strategy here is to require that the final fitted values (posterior parameter estimates) should remain relatively close to the anatomical connectivity priors, and retain the overall characteristics of that matrix in terms of weight distributions and other spatial features. To ensure the posterior connection weights remain close to the anatomical priors, we employ a regularization approach that normalize the connectivity weights matrix by dividing it by its norm, effectively constraining the maximum eigenvalue of the connectivity matrix to 1. This approach ensures that while individual connection strengths may vary during optimization, the overall connectivity structure remains consistent with known anatomical constraints. Importantly, after fitting the model to each subject's data, we conducted a visual inspection to verify that the posterior mean connection weight matrices preserved the essential topological features observed in empirical neuroimaging data.

Similarly, the projection matrix from sources to channels, computed by solving the Maxwell equations with a boundary element method[34] using MNE[35], begins with a set of population-representative priors, and is allowed to vary per subject with tight constraints by setting small prior variance. For a comprehensive overview of the distribution of the optimized model parameters, please consult Fig. S1.

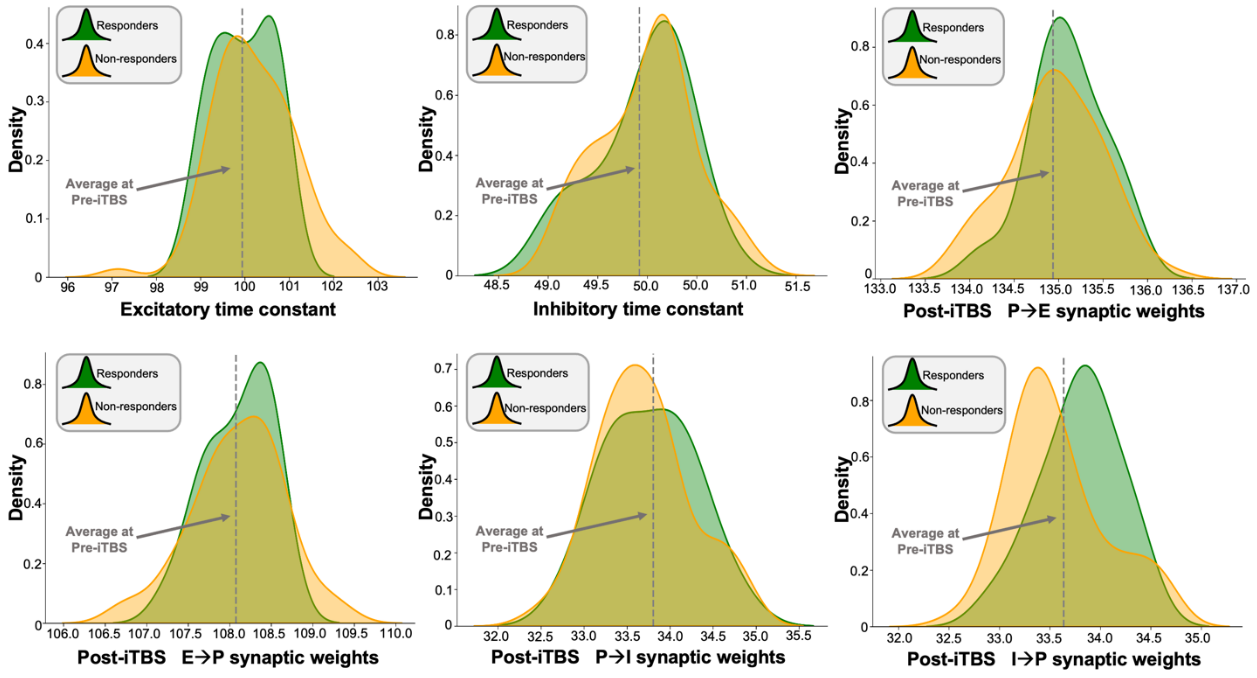


**Fig. S1. Distributions of physiological parameter estimates for responders and non-responders.** Density plots showing the optimized values for the Jansen-Rit model's physiological parameters across responders (green) and non-responders (yellow) post-iTBS treatment. The gray dashed line indicates the average value pre-iTBS for all subjects. Parameter estimation was performed using our novel automatic differentiation and gradient-based approach inspired by current techniques in deep learning[2] that we also developed in our other studies[2,17].

### 2. SUPPLEMENTARY RESULTS

#### 2.1 Non-significant Effects of iTBS on Physiological Parameters

While our main manuscript reports significant effects for inhibitory-to-pyramidal connectivity (I→P), we conducted similar analyses for all other physiological parameters in the Jansen-Rit model. The TIME×GROUP interaction was non-significant for excitatory time constant ($F(1,86) = 2.37$, $p = 0.127$), inhibitory time constant ($F(1,86) = 1.82$, $p = 0.181$), pyramidal-to-excitatory connectivity (P→E; $F(1,86) = 1.94$, $p = 0.168$), excitatory-to-pyramidal connectivity (E→P; $F(1,86) = 1.73$, $p = 0.192$), and pyramidal-to-inhibitory connectivity (P→I; $F(1,86) = 0.98$, $p = 0.326$).

These findings indicate that the I→P connectivity parameter was uniquely affected by iTBS treatment in responders compared to non-responders, while other physiological parameters of the Jansen-Rit model remained statistically unchanged between groups. This suggests a selective mechanism of action for effective iTBS intervention that primarily involves modulation of inhibitory influence on pyramidal cells.

#### 2.2 Connectome-based neurophysiological modelling accurately reproduces subject-specific TMS-EEG dynamics

As an important preliminary result, extensive testing confirmed that our new connectome-based neurophysiological model of stimulus-evoked responses achieves robust and accurate recovery of measured TMS-EEG time series at both the group-average and individual-subject levels. Fig. S2A shows empirical and fitted (i.e. simulated, with optimized physiological parameters) TMS-EEG waveforms along with selected topography maps for two example subjects. It is visually evident in these figures that the model accurately captures several individually-varying features of the TMS-EEG time series, such as the timing of the early and late components, and the extent to which they are dominated by left/right and temporal/parietal/frontal channels. For the latter, this can be seen by comparing the line colors in the upper and lower rows of corresponding columns in Fig. S2, and using the channel location references given by the channel color map on the top left of each topoplot. Pearson correlations between empirical and simulated TMS-EEG time series confirmed that an excellent goodness-of-fit was observed at the individual channel level. Time-wise permutation tests revealed a significant Pearson correlation coefficient for every electrode, with an average $R^2$ value of 93% across all subjects ($p < 0.0001$). As well as the millisecond-by-millisecond comparisons and the timing of key waveform components, we also assessed the accuracy of the model in capturing holistic time series properties for both early and late responses.

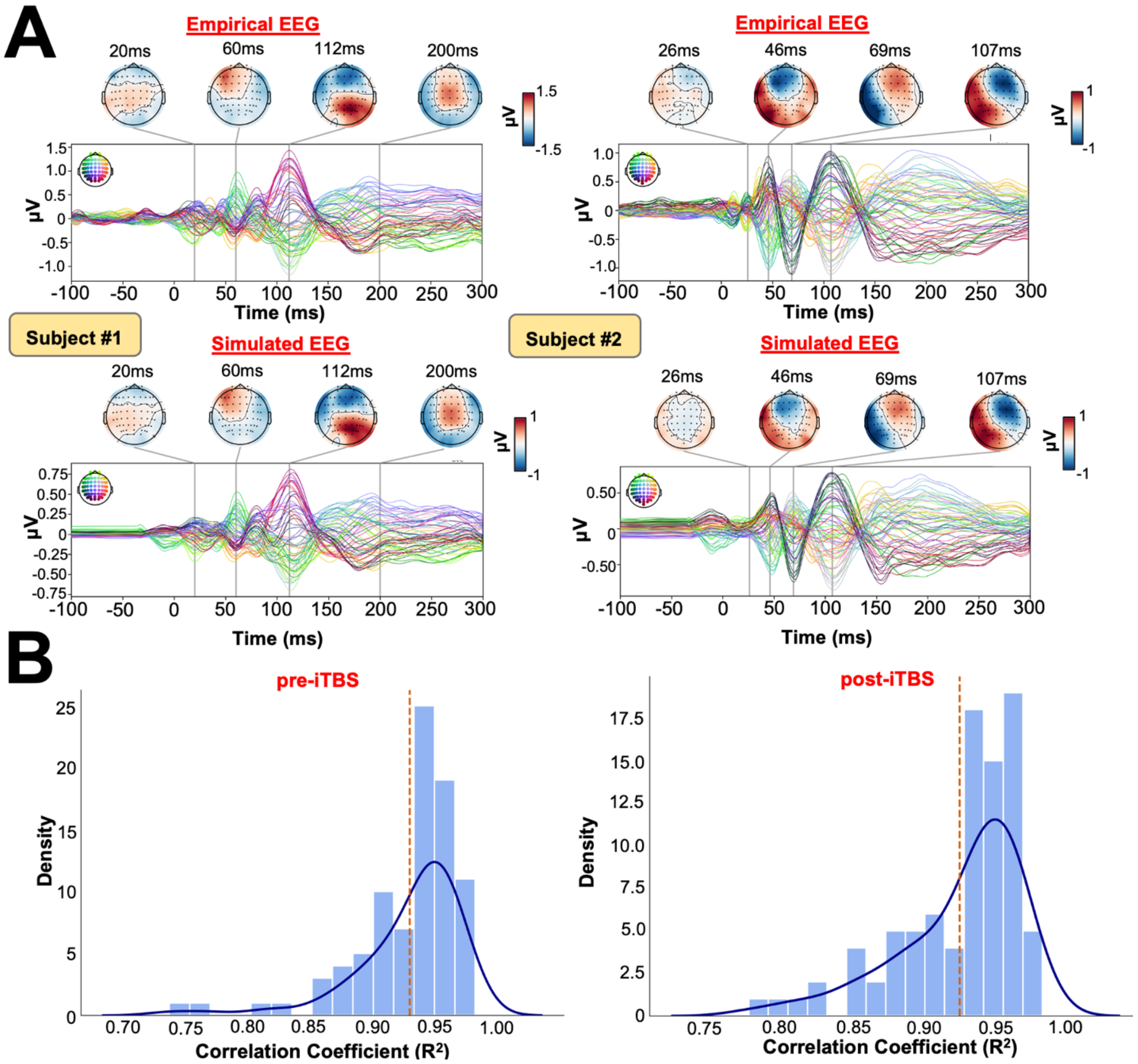


**Fig. S2. Goodness-of-fit between empirical and simulated TMS-EEG data. (A)** Empirical (upper row) and simulated (lower row) TMS-EEG butterfly plots, with scalp topographies for three representative subjects. These simulations, showing a robust recovery of individual empirical evoked potential patterns in model-generated activity EEG time series. **(B)** Histogram plots showing the distributions of the Pearson correlation coefficients between simulated and empirical time series for each subject and session. The left panel shows pre-iTBS data, and the right panel shows post-iTBS data. The dashed red line indicates the average Pearson correlation across subjects.

#### 2.3 TMS-Evoked Spectral Power in Responders and Non-responders

We examined the TMS-evoked spectral power patterns in both responders and non-responders before and after iTBS treatment. As shown in Fig. S3, time-frequency representations of the power differences (post-iTBS minus pre-iTBS) revealed distinct patterns between groups. Time-frequency analysis revealed distinct patterns between groups, with responders showing a more pronounced reduction in low-frequency (3-10 Hz) power following iTBS compared to non-responders. This power suppression was particularly evident in the 50-200ms post-stimulus window. While both groups exhibited some degree of post-treatment change in induced spectral power, the magnitude and spectral-temporal extent of these changes differed substantially. Non-responders showed a more limited pattern of spectral modulation mainly in the higher frequency bands, whereas responders demonstrated a broader suppression effect spanning theta-alpha frequencies. These group-level power spectra provided the basis for the second-level statistical analysis presented in the main text, which quantified the significant between-group differences in treatment-induced spectral modulation.

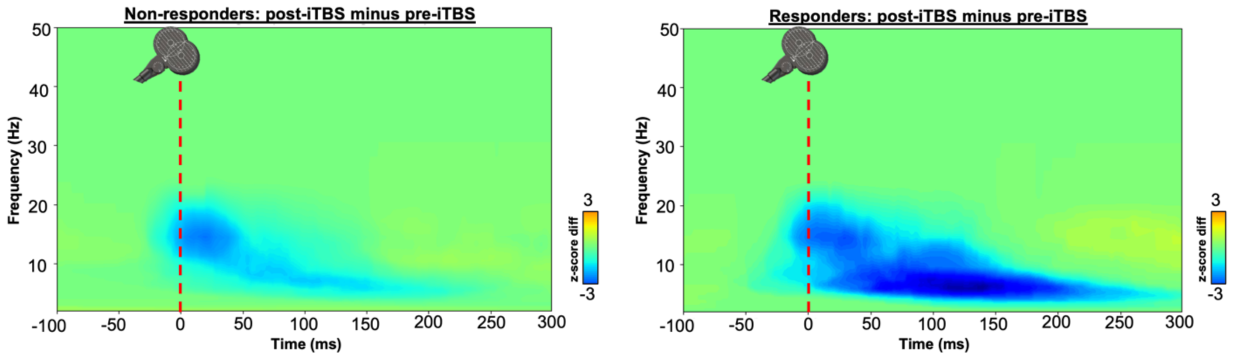


**Fig. S3. Group-level TMS-induced spectral power changes following iTBS treatment. (A)** Time-frequency representation of TMS-evoked power differences (post-iTBS minus pre-iTBS) for non-responders. **(B)** Time-frequency representation for responders. Color scale represents z-score normalized power changes, with blue indicating post-treatment power reduction. Vertical red dashed line indicates TMS pulse onset. Note the more pronounced low-frequency (3-10 Hz) power suppression in responders between 50-200ms post-stimulus compared to non-responders. These first-level spectral maps formed the basis for the statistical comparisons reported in Fig. 1A.

### 3. SUPPLEMENTARY VIDEOS

**Video V1. Dynamic visualization of TMS-EEG neural trajectories pre-iTBS.** Temporal evolution of neural trajectories in the treatment-predictive state-space, highlighting responders (green) and non-responders (orange) before iTBS. Prior to treatment, both groups exhibit highly similar trajectories and remain close to the stable fixed point (black dot), indicating no substantial differences in the underlying dynamical regime.

**Video V2. Dynamic visualization of TMS-EEG neural trajectories post-iTBS.** Temporal evolution of neural trajectories in the treatment-predictive state-space, showing post-treatment dynamics for responders (green) and non-responders (orange). Post-iTBS, responders exhibit a marked contraction of the trajectory and reduced deviation from the fixed point, indicating increased dynamical stability and reduced sensitivity to external perturbations such as TMS.
